# CCR3 inhibition suppresses inflammation-driven recruitment of peripheral immune cells to the eye

**DOI:** 10.1101/2023.09.08.556878

**Authors:** Yesenia Lopez, Sofia Caryotakis, Sharda Raina, Sanket V. Rege, Reema Harish, Rebecca Ray, Idit Kosti, Arnaud Teichert, S. Sakura Minami, Onkar S. Dhande

## Abstract

C-C chemokine receptor type 3 (CCR3) has been linked with age-related macular degeneration (AMD) pathologies. Specifically, its function as an immune modulator in AMD remains unclear. To address this question, we investigated the impact of CCR3 inhibition on inflammation, a key driver of AMD pathologies, by assessing inflammatory cytokines and infiltrating immune cells in two models of ocular inflammation.

Mice were orally dosed twice a day with AKST4290, a CCR3 small molecular inhibitor, in the sodium iodate (NaIO3) and myelin oligodendrocyte glycoprotein (MOG) models. A combination of autoradiography and analytical chemistry techniques were used to assess drug concentration and distribution. Bead-based multiplexing technology was used to determine cytokine concentrations, and flow cytometry and immunohistochemistry was used to ascertain ocular immune-cell composition.

CCR3 expression was detected in the retinal pigment epithelium (RPE)/choroid complex where AKST4290 was found to preferentially accumulate at sustained levels. In the NaIO_3_ model, inhibition of CCR3 with AKST4290 significantly decreased both the concentration of specific chemokines and the number of multiple populations of infiltrating peripheral immune cell. Furthermore, effects of CCR3 inhibition on immune cell infiltration were confirmed in the MOG model.

These data demonstrate that CCR3 inhibition strongly modulates local inflammation by impacting both cytokine concentrations and immune cell composition in ocular diseases.

Moreover, these findings together with the known role of CCR3 in promoting pathologic angiogenesis implicate a pleiotropic role for CCR3 in AMD.

## Introduction

Inflammation plays a central role in the pathogenesis of multiple ocular diseases including AMD [1,2]. In early-stage AMD, RPE cells release multiple chemokines that serve as recruitment signals for immune cells [3,4]. These infiltrating immune cells further exacerbate the inflammatory milieu by releasing pro-inflammatory factors that in turn promote recruitment of more immune cells into the eye. Unchecked this feedback loop can drive dysfunctional and chronic inflammation [5]. In both dry and neovascular AMD (nAMD), such runaway inflammation is thought to promote RPE degeneration, drusogenesis, vascular leakage, and choroidal neovascularization (CNV) and these pathologies ultimately lead to blindness [6–8].

Current FDA approved therapies for nAMD almost exclusively target the blockade of pro-angiogenic vascular endothelial growth factor (VEGF). Additionally, only recently has the first FDA approved therapeutic for dry AMD, targeting complement C3 inhibition, come to market [9]. In both cases, AMD treatments require repeated intravitreal injections which is not only a burden on patients and clinicians, but also carries the risk of infection and complications such as ocular hemorrhaging [10]. Moreover, several complement and VEGF-independent pathways are implicated in driving inflammation in dry and nAMD respectively and are being actively explored as therapeutic targets [11,12].Therefore, an oral self-administered drug targeting novel pathways presents a compelling entry point for AMD therapeutic development.

In recent Phase 2a clinical studies, inhibition of CCR3 with AKST4290 was associated with stable or improved best-corrected visual acuity after 6 weeks in the majority of both treatment-naïve and refractory nAMD subjects [13]. CCR3 is a G-protein coupled receptor that is involved in multiple functions including angiogenesis, immune cell recruitment into sites of inflammation and allergic responses [14–16]. While the role of CCR3 signaling in neovascularization is well established its impact on ocular inflammation, a key factor in the development and progression of AMD, remains unclear [3,17,18]. CCR3 ligands, notably CCL11 (eotaxin), can act as proinflammatory immune signals [19]. CCL11 is increased in human and mouse plasma with aging and linked to inflammation-driven dysfunction in the CNS [20,21].

To elucidate the role of CCR3 in modulating inflammation in the eye, we investigated the effects of CCR3 inhibition in two mouse models of ocular inflammation: sodium iodate (NaIO_3_) and myelin oligodendrocyte glycoprotein (MOG) [22,23]. First, we used a combination of autoradiography and analytical chemistry techniques to assess AKST4290 concentration and distribution in the mouse eye. Next, using a combination of bead-based multiplexing technology, flow cytometry, and immunohistochemistry we examined the effects of CCR3 inhibition by AKST4290 on inflammatory cytokines and infiltrating immune cells. Together these data shed new light on the function of CCR3 in inflammation and disease and advances the mechanism of AKST4290 in the eye.

## Methods

### Animals

Mice were purchased from Hilltop Lab Animals (Scottsdale, PA) and The Jackson Laboratory (Bar Harbor, ME, USA) and treated in accordance with IACUC policy and the ARVO Statement for the Use of Animals in Ophthalmic and Vision Research. All animals were housed at a standard temperature (22 ± 1°C) in a light-controlled environment (light-cycle from 7 am to 7 pm) with *ad libitum* access to food and water.

### Quantitative whole-body autoradiography (QWBA)

QWBA was performed and analyzed by WuXi AppTec (Cranbury, NJ). A stock [^14^C]- AKST4290 solution was formulated with ethanol. This stock was then mixed with AKST4290 and purified water to obtain a 1.25 mg/mL dosing solution that was administered orally (single, 10 mg/kg, 100 µCi) to 6 week-old C57BL/6 mice. Autoradiography was performed at various post-dose time points [15 mins to 2016 hours (12 weeks)] to visualize drug distribution in the uveal tract, lens, and eye (whole eye minus uveal tract and lens). In brief, carcasses were embedded in low viscosity carboxymethylcellulose and sagittal 30 µm sections of the whole body were sectioned using a Leica 9800 Cryomicrotome (Leica Biosystems, St. Louis, MO). Whole body sections were placed in direct contact with phosphor imaging plates (BAS-SR 2025, Fujifilm, Tokyo, Japan) for approximately 4 days at room temperature. Finally, autoradioluminograms were generated with a GE Typhoon FLA 9500 Phosphor Imager (Pittsburgh, PA) and analyzed using AIDA software (Raytest GmbH, Berlin, Germany). Concentration was calculated as:

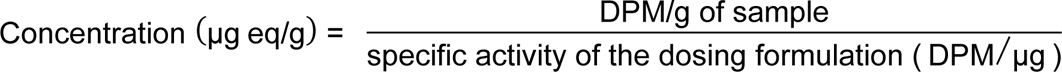

*DPM = disintegrations per minute

### Liquid chromatography with tandem mass spectrometry (LC-MS/MS)

LC-MS/MS was performed and quantified by Quintara Discovery (Hayward, CA). AKST4290 was formulated in HP-β-cyclodextrin (Sigma-Aldrich, St. Louis, MO) to 9 mg/mL, adjusted to pH 6.5, and a single dose of 30 mg/kg was administered orally to 8 week-old C57BL/6 mice. At 0.5, 2, and 24 hours post-dose, mice were deeply anesthetized with Avertin (Sigma-Aldrich, St. Louis, MO), saline perfused, and eyes were harvested at each timepoint. The eyes were dissected in ice-cold 1x PBS (phosphate buffered saline) to obtain the retina and the RPE/choroid/sclera complex, which were subsequently snap frozen. Frozen tissue was homogenized with three volumes of ice-cold water and the homogenate was then extracted with actenitrile containing internal standard (50 ng/mL of dexamethasone). The subsequent supernatant was used for LC-MS/MS on a Sciex Qtrap 6500+ (Framingham, MA). PK parameters were determined via noncompartmental analysis (NCA) with Phoenix 64 WinNonLin (Certara, Princeton, NJ).

### Sodium iodate (NaIO_3_) model induction and dosing paradigm

12 to 14 week-old C57BL/6 mice were divided into three groups: Vehicle, NaIO_3_ only, and NaIO_3_ + AKST4290. AKST4290 treated mice received 60mg/kg doses, while the Vehicle and NaIO_3_ only groups received vehicle solution without AKST4290, HP-β- cyclodextrin (Sigma-Aldrich, St. Louis, MO) and H_2_O (3:1 volume ratio) adjusted to pH 6.5 with 1M NaOH. Both vehicle and AKST4290 were administered via oral gavage (PO) twice daily (BID) approximately 8 hours apart. NaIO_3_ (20 mg/kg) or PBS (Vehicle group only) was injected intravenously (IV; tail vein) ∼1-2 hours following the first AKST4290 or vehicle solution PO dose. The last dose of AKST4290 or vehicle solution was administered 3-4 hours before sacrifice on day 3.

### Ocular histology

#### RPE flatmounts

Eyes were harvested from saline perfused mice and fixed in 4% PFA (paraformaldehyde) overnight at 4°C. RPE flatmounts were stained with Alexa Fluor plus 647-Phalloidin (1:10,000, Thermo Fisher, Waltham, MA). Confocal z-stack images of phalloidin stained RPE flatmounts were acquired using the Zeiss LSM800 (White Plains, NY). To quantify the area of damage, maximum intensity projections were generated and average radius ratio, defined as the average of the (length of damage) / (length of tissue) for each leaflet in the RPE flatmount, was computed using ImagePro (Media Cybernetics, Rockville, MD). Only mice with average radius ratio greater than 0.5 (i.e. >50% area of damage), were included in subsequent analyses.

#### Retinal flatmounts

Eyes were harvested from saline perfused mice and fixed in 4% PFA for 2-4 hours at 4°C. Retinal flatmounts were stained with CD3 (1:100, BD Biosciences, Franklin Lakes, NJ) and Collagen IV (1:50, EMD Millipore, Burlington, MA) for 48 hours at 4°C and then secondary antibodies donkey anti-rat Alexa Fluor 488 and donkey anti-goat Alexa Fluor 555 (1:300, Invitrogen, Waltham, MA) at room temperature for 2 hours. Total retinal CD3+ cells per retina were counted live using the Zeiss LSM800 (White Plains, NY).

### Ocular cytokine analysis

Eyes were harvested from saline perfused mice and snap frozen. Lysates were prepared by homogenizing a whole frozen saline perfused eye in RIPA (EMD Millipore, Burlington, MA) + protease inhibitor (Thermo Fisher, Waltham, MA) with a bead ruptor (Omni International, Kennesaw, GA). Lysates were centrifuged for 10 mins at 10,000 g at 4°C to collect the supernatant. Protein concentration was determined by BCA (protein assay, Thermo Fisher, Waltham, MA), and supernatant was diluted with DPBS (Gibco, Grand Island, NY) to 1 mg/mL accordingly. Samples were analyzed by Eve Technologies (Calgary, Canada) using the Mouse Cytokine/Chemokine 31-Plex Discovery Assay Array.

### Flow cytometry

Mice were deeply anesthetized with isoflurane followed by euthanasia by cervical dislocation. Eyes were harvested, the RPE/choroid/sclera complex were dissected and subsequently digested at 37°C for 45 min using the Multi Tissue Dissociation Kit 1 (Miltenyi, Bergisch Gladbach, Germany). 1 mL of enzyme solution was used per sample. Softened fragments of tissue were gently triturated then pushed through a 70 µm nylon cell strainer (Corning, Corning, NY). Next, single-cell suspensions were immunostained for cell surface markers for various immune cell types. Zombie Red viability dye was used to distinguish live and dead cells (Biolegend, San Diego, CA). Fc receptors were blocked with anti-CD16/32 in the antibody staining solutions (BD Biosciences, Franklin Lakes, NJ). For RPE and endothelial identification, cells were stained with PerCP-labeled CD45, PE-Cy7-labeled CD31 (endothelial marker), Alexa Fluor 488-labeled CD140b (RPE marker, Plaza Reyes et al., 2020), and PE-labeled CCR3 (CD45 & CD31: Biolegend, San Diego, CA; CD140b: Thermo Fisher, Waltham, MA; CCR3: Novus Biologicals, Centennial, CO). For myeloid cell lineages, cells were stained with B711-labeled mouse CD11b, PB-labeled Ly6G, BV510-labeled Ly6C, PE-labeled CD206, PE-Cy7-labeled CX3CR1, and FITC-labeled CD45 (Biolegend, San Diego, CA). For T cell analysis, cells were stained with eFluoro780-labeled CD3, eFluoro506-labeled CD4, and SuperBright 600-labeled CD8 (Biolegend, San Diego, CA). Immunostained cells were analyzed using an Attune NxT flow cytometer (violet, blue, yellow, and red lasers configuration, Thermo Fisher, Waltham, MA).

### Myelin oligodendrocyte glycoprotein (MOG) model and dosing paradigm

Male 9 week-old C57BL/6 mice were divided into three groups (Vehicle, MOG only, MOG + AKST4290) and dosed with either AKST4290 (30 mg/kg) or vehicle solution, HP-β-cyclodextrin (Sigma-Aldrich, St. Louis, MO) and H_2_O adjusted to pH 6.5 with 1M NaOH, via oral gavage, twice daily for 9 days, starting 2 hours after MOG induction. MOG was induced with 100 µL SQ injections in each flank of emulsified myelin oligodendrocyte glycoprotein, MOG35-55 4mg/mL in PBS (Anaspec, Fremont, CA), and M. Tuberculosis H37 Ra (Fisher Scientific, Waltham, MA) dissolved in complete Freud’s adjuvant, CFA. Pertussis toxin was IP administered on the day of induction (0.002 mg/mL) and again 2 days later.

### Statistical analysis

Statistical analysis was performed in Prism 9 (GraphPad, San Diego, CA) and R Statistical Software (version 4.2.0; R Foundation for Statistical Computing, Vienna, Austria). For cytokine and immune cell analyses, a nested linear model followed by Tukey’s posthoc analysis was used to assess statistical significance. For all other analyses, a Mann-Whitney test was used. p-value significance and n values for each experiment are indicated in the figure legends. The n values represent biological replicates.

## Results

### CCR3 is expressed by ocular cell-types affected by AMD

The choroid and RPE are key sites of pathology in AMD and are comprised primarily of endothelial and RPE cells. To determine if CCR3 is expressed by these cell types, flow cytometry was performed on isolated RPE/choroid complexes. Single-cell suspensions were generated from individual healthy RPE/choroid complexes and immunolabelled them with a cocktail of antibodies (see methods for details) to probe for CCR3 expression on both endothelial and RPE cells (Fig 1A). A substantial population of RPE (14.68% ± 0.83, n = 5) and choroidal endothelial cells (18.14% ± 1.07, n = 5) do indeed express CCR3 (Fig 1B).

**Fig 1.**
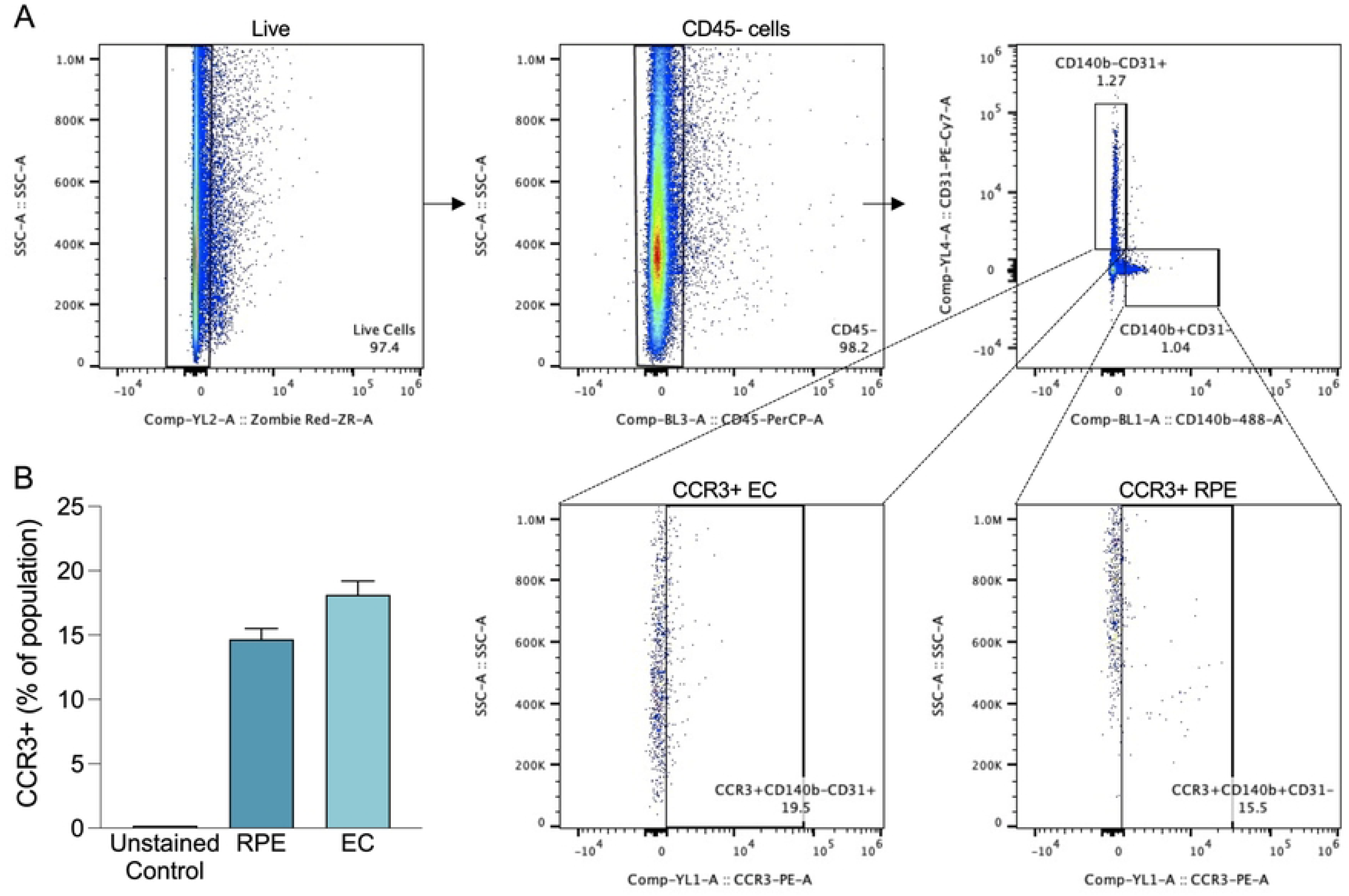
Ocular cell-types impacted by AMD express CCR3. (A) Gating strategy for identifying cell types and CCR3 expression using flow cytometry. (B) Proportion of RPE and EC expressing CCR3 (n=5). Mean ± SEM. EC: endothelial cells; RPE: retinal pigment epithelial cells.

### AKST4290 preferentially accumulates at the site of AMD pathology

Next, we sought to define the ocular levels and localization of AKST4290, a small molecule inhibitor of CCR3. First, QWBA was performed to determine the dynamic accumulation and metabolism of radioactively labelled AKST4290 after a single oral dose in mice (Fig 2A). Based on the limited resolution of QWBA image data, two broad subdivisions of the eye, uveal tract and the rest of the ocular structure termed eye were defined. Measurable levels of [^14^C]-AKST4290 were detected as early as 15 minutes post-dose (eye: 0.12 µg eq/g, uveal tract: 0.27 µg eq/g, n =1). AKST4290 levels peaked in both structures by 24 hours post-dose (eye: 0.56 µg eq/g, uveal tract: 2.61 µg eq/g, n=1) and were maintained up to 12 weeks post-dose in the uveal tract (0.18 µg eq/g, n =1) of male C57BL/6 mice (Fig 2B). Similarly, in female mice AKST4290 levels peaked in both structures at the same time, 48 hours, (eye: 0.3 µg eq/g, uveal tract: 1.37 µg eq/g, n=1) and sustained in the uveal tract 12 weeks post-dose (0.06 µg eq/g, n=1; Fig 2C). Next, to ascertain the levels of AKST4290 more precisely within the RPE/choroid complex and retina, LC-MS/MS was performed on sub-dissected ocular tissues collected after a single oral dose of AKST4290. At 30 minutes post-dose, AKST4290 concentrations were substantially higher in the RPE/choroid complex in comparison to the retina (RPE/choroid: 447.8 ng/mL ± 143.89, retina: 26.11 ng/mL ± 11.15). Moreover, these levels were sustained up to 24 hours within both structures (RPE/choroid: 941.5 ng/mL ± 273.09, retina: 113.05 ng/mL ± 65.03; Fig 2D).

**Fig 2.**
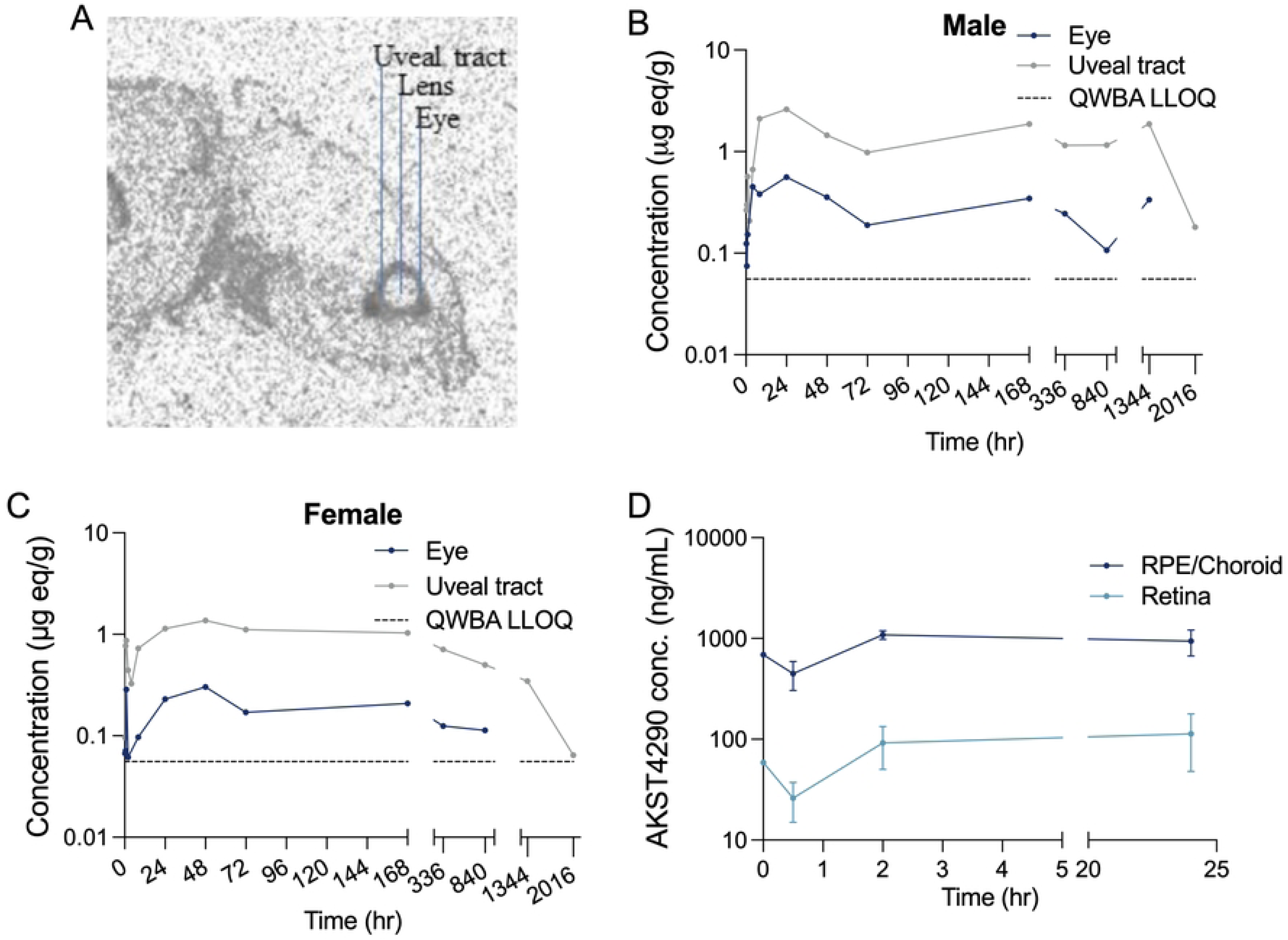
AKST4290 levels in the eye after a single dose. (A) Representative section of QWBA highlighting eye compartments. (B,C) Quantification of [14C]-AKST4290 levels after a single PO dose of 10mg (100 µCi/kg) in male (B; n=1/timepoint) and female (C; n=1/timepoint) mice. (D) Quantification of AKST4290 levels in eye sub compartments after a single 30 mg/kg PO dose (n=1 at timepoint 0, n=4-5/all other timepoints). Mean ± SEM. PO.: oral gavage; QWBA: quantitative whole-body autoradiography; LLOQ: lower limit of quantification.

### CCR3 inhibition reduces inflammatory chemokines in the NaIO_3_ model

To investigate the impact of CCR3 inhibition on ocular inflammation, we utilized the NaIO_3_ model. Initially, the model was established and confirmed by assessing RPE damage after a single IV injection of PBS or NaIO_3_ (20mg/kg) at 3 days post injection. This confirmed that control PBS injection did not cause damage to the RPE as evidenced by the intact and uniform phalloidin labeling of RPE cells visualized in a flatmount preparation (Fig 3A, left panel). In contrast, NaIO_3_ (20mg/kg) caused robust central-to-peripheral degeneration of the RPE layer at the same time point as revealed by a striking absence of phalloidin labeled RPE cells (Fig 3A, right panel and S1 Fig). Next, the impact of CCR3 inhibition on ocular inflammation was assessed at 3 days post PBS or NaIO_3_ (20mg/kg) injection with an interventional dosing paradigm of daily twice a day dosing (PO BID) of AKST4290 (60mg/kg) or vehicle solution (Fig 3B). NaIO_3_ model induction in each mouse was confirmed by computing the extent of RPE degeneration in phalloidin stained RPE flatmount of one eye. PBS injected mice treated with vehicle solution did not impact the RPE layer (0%; n=16), whereas comparable levels of NaIO_3_-induced RPE loss was observed between the NaIO_3_ (82.34 ± 1.41%; n=16) and the NaIO_3_ + AKST4290 (82.22 ± 0.85%; n=16; p=0.724) treated mice (Fig 3C). One measure of ocular inflammation is the level of inflammatory cytokines present in ocular tissue and fluids. The concentration of several inflammatory cytokines in whole eye homogenates was quantitatively assessed (Fig 3 and S2 Fig). CCL11, the primary ligand of CCR3, was significantly increased by NaIO_3_-induced damage (p < 0.001, Veh: 6.6 ± 0.3 pg/mL, n = 25, NaIO_3_: 8.8 ± 0.33 pg/mL, n = 23) and remained unchanged with AKST4290 dosing (p = 0.367, NaIO_3_ + AKST4290: 8.2 ± 0.24 pg/mL, n = 30; Fig 3D). In addition, VEGF, a key driver of neovascular AMD pathologies, was significantly decreased in the NaIO_3_ (p < 0.001, Veh: 1.7 ± 0.14 pg/mL, n = 25, NaIO_3_: 0.8 ± 0.07 pg/mL, n = 23) model (Fig 3E). Excitingly, CCR3 inhibition had a major impact on NaIO_3_ induced increases in pro-inflammatory chemokines CCL5 and CXCL9. Both CCL5 (Veh: 1.41 ± 0.1 pg/mL, n = 22, NaIO_3_: 2.05 ± 0.13 pg/mL, n = 24; p < 0.01) and CXCL9 (Veh: 1.8 ± 0.26 pg/mL, n = 24, NaIO_3_: 5.6 ± 0.67 pg/mL, n =25; p < 0.001) were elevated with NaIO_3_ injury and significantly decreased with AKST4290 administration (CCL5: NaIO_3_ + AKST4290: 1.8 ± 0.12 pg/mL, n = 29, p < 0.01; CXCL9:NaIO_3_ + AKST4290: 3.7 ± 0.41 pg/mL, n = 29; p < 0.05; Figs 3F and 3G).

**Fig 3.**
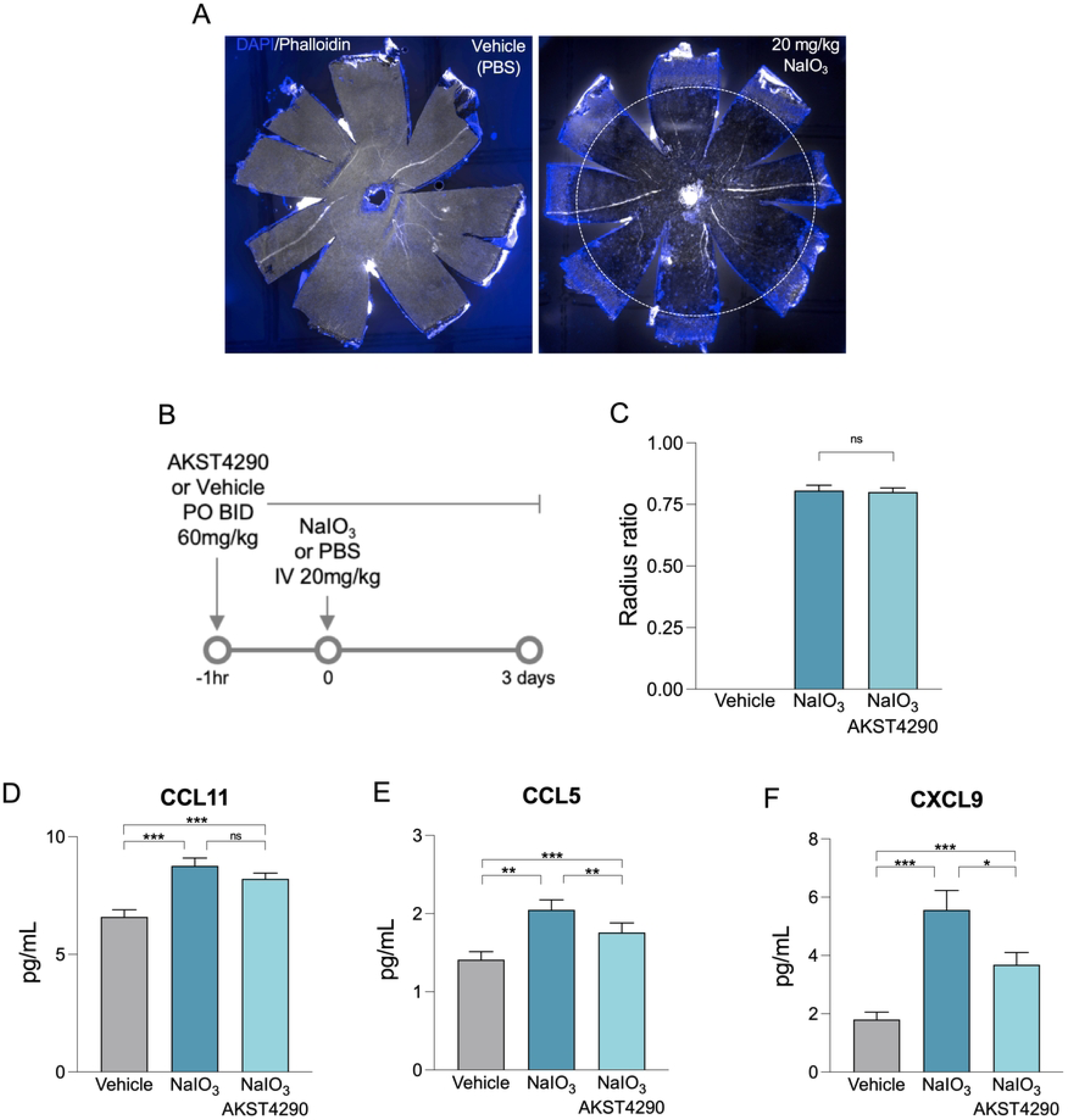
CCR3 inhibition reduces inflammatory chemokines in the NaIO_3_ model. (A) RPE flatmount stained with phalloidin (white) and DAPI (blue) showing RPE degeneration (dotted circle) at 3 days post single IV injection of 20mg/kg NaIO_3_ (right panel) or vehicle (left panel). (B) NaIO_3_ model and AKST4290 dosing paradigm. (C) Quantification of radius ratio (average radial RPE damage) in RPE flatmounts of mice treated with vehicle, NaIO_3_, or NaIO_3_ + AKST4290. (D-F) Chemokine levels of CCL11 (D), CCL5 (E), and CXCL9 (F). Mean ± SEM. n=22-30 mice/group (C-F). *p<0.05, **p<0.01, ***p<0.001, ****p<0.0001; nested linear model followed by Tukey’s posthoc analysis. ns: not significant; PO: oral gavage; BID: twice daily; IV: intravenous.

### CCR3 inhibition reduces immune cell infiltration into the RPE

To further elucidate the impact of CCR3 inhibition on ocular inflammation flow cytometry was used to quantitively assess effects of AKST4290 on immune cell infiltration in the RPE/choroid complex, the primary site of NaIO_3_ injury. Single-cell suspensions of the RPE/choroid complex were generated, labeled with immune cell-type specific markers, and gated to delineate different myeloid and lymphocyte populations (Fig 4A). NaIO_3_ injury resulted in a significant increase in infiltrating myeloid cells in comparison to age-matched controls (Veh: 1866 ± 326, NaIO_3_: 7046 ± 802, n = 12/group, p < 0.0001; Fig 4B). This increase in infiltrating myeloid cells was significantly reduced following CCR3 inhibition by AKST4290 (NaIO_3_ + AKST4290: 4905 ± 619, n = 12; p = 0.0007). Furthermore, AKST4290 dosing significantly decreased NaIO_3_-induced infiltration of two myeloid derived populations namely neutrophils (Veh: 14 ± 3, NaIO_3_: 146 ± 17, NaIO_3_ + AKST4290: 82 ± 15, Vehicle vs. NaIO_3_: p < 0.0001, NaIO_3_ vs. NaIO_3_ + AKST4290: p = 0.002; Fig 4C) and monocytes (Veh: 74 ± 16, NaIO_3_: 1679 ± 222, NaIO_3_ + AKST4290: 1051 ± 153; Vehicle vs. NaIO_3_: p < 0.0001, NaIO_3_ vs. NaIO_3_ + AKST4290: p = 0.0036; Fig 4D). Although not significant, inhibition of CCR3 by AKST4290 trended to reduce the numbers of infiltrating macrophages resultant from NaIO_3_-mediated injury (Veh: 350 ± 90, NaIO_3_: 656 ± 131, NaIO_3_ + AKST4290: 484 ± 94, Vehicle vs. NaIO_3:_ p = 0.0015; NaIO_3_ vs. NaIO_3_ + AKST4290 p = 0.0941; Fig 4E). Next, the effect of CCR3 inhibition on infiltrating lymphocyte populations was examined. NaIO_3_ resulted in a significant increase in CD3+ lymphocytes in the RPE/choroid and no change was observed with AKST4290 administration (Veh: 77 ± 12, NaIO_3_: 305 ± 35, NaIO_3_ + AKAT4290: 225 ± 43, Veh vs. NaIO_3_ p < 0.0001, NaIO_3_ vs. NaIO_3_ + AKST4290 p = 0.2189; Fig 4F). Interestingly, when probed for specific types of CD3+ T cells it was discovered that although inhibition of CCR3 with AKST4290 did not reduce CD8+ lymphocytes (Veh: 2 ± 1, NaIO_3_: 29 ± 7, NaIO_3_ + AKST4290: 21 ± 4, Vehicle vs. NaIO_3_: p < 0.0001, NaIO_3_ vs. NaIO_3_ + AKST4290 p = 0.2295; Fig 4G), inhibition of CCR3 with AKST4290 did significantly reduce infiltrating CD4+ lymphocytes (Veh: 33 ± 5, NaIO_3_: 170 ± 26, NaIO_3_ + AKST4290: 131 ± 29, Veh vs. NaIO_3_: p < 0.0001, NaIO_3_ vs. NaIO_3_ + AKST4290: p < 0.0001; Fig 4H).

**Fig 4.**
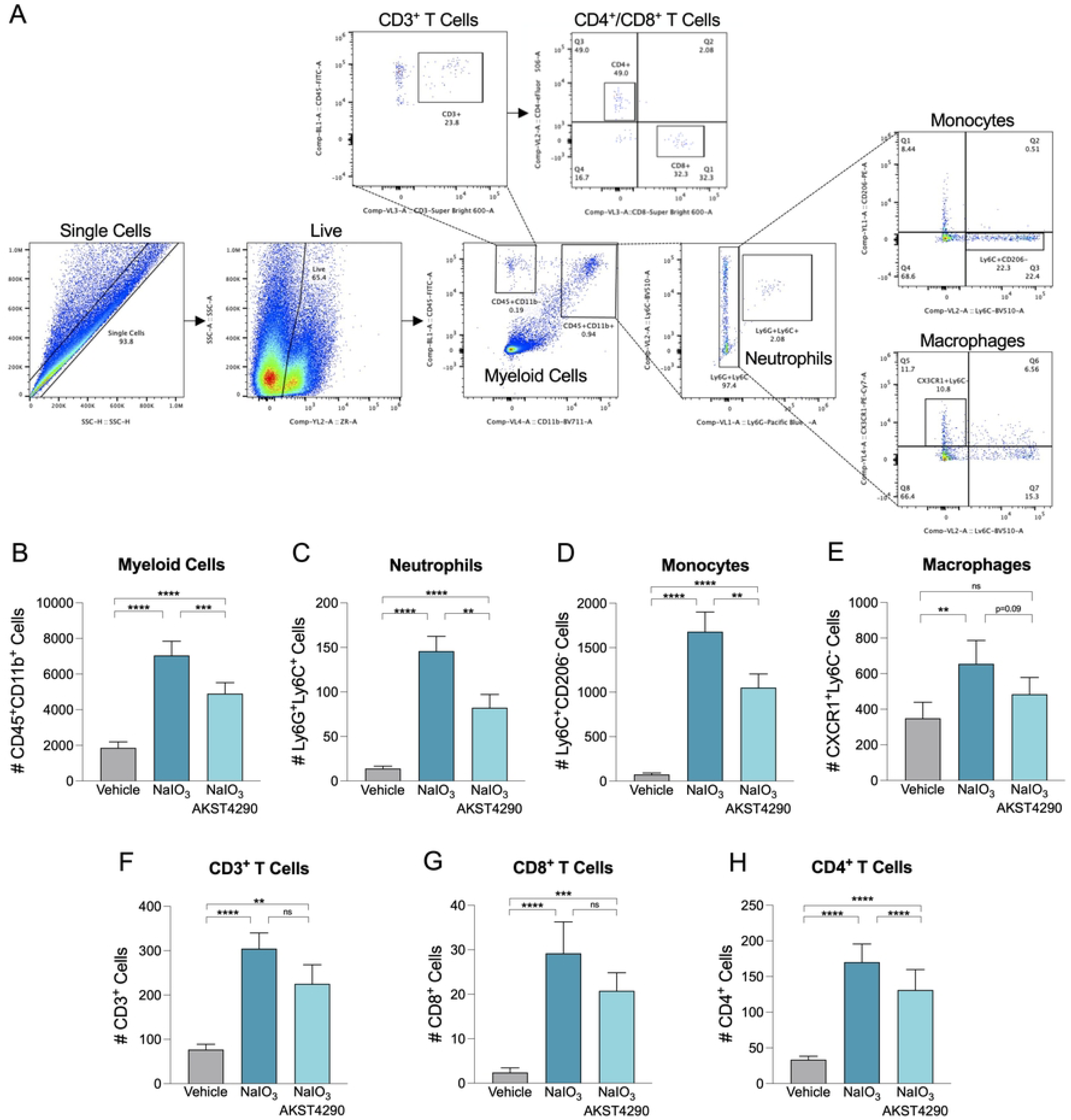
CCR3 inhibition reduces NaIO_3_-induced immune cell infiltration into the RPE/choroid complex. (A) Gating strategy for identifying various immune cell populations using flow cytometry. Quantification of myeloid cells (CD45+CD11b+) (B), neutrophils (Ly6G+Ly6C+) (C), monocytes (Ly6C+CD206-) (D), microglia/macrophages (CX3CR1+Ly6C-) (E), and CD3+ (F), CD4+ (G), and CD8+ (H) T cells in mice treated with Vehicle, NaIO_3_, or NaIO_3_ + AKST4290. Mean ± SEM. **p<0.01, ***p<0.001, ****p<0.0001; nested linear model followed by Tukey’s posthoc analysis. ns: non-significant. n=12 mice/group (B-H).

### CCR3 inhibition reduces retinal T cell infiltration in a model of induced autoimmune inflammatory disease

To specifically assess T cell infiltration in the retina, a distinct facet of AMD pathology, a model of autoimmune inflammatory disease was utilized. An immune reaction was generated against myelin with M. Tuberculosis H37 Ra and pertussis toxin in a similar fashion to experimental autoimmune encephalomyelitis (EAE) induction [25]. On an abbreviated time course, in comparison to EAE, this technique has been shown to robustly mobilize T cells to myelin containing regions such as the CNS and we refer to this model as MOG (myelin oligodendrocyte glycoprotein) [23]. Thus the MOG model does not recapitulate AMD but rather allows for validation of the robust effect CCR3 inhibition has on a specific feature of the immune response in AMD. At 9 days after MOG induction, T cells were clearly observable in the retinal vasculature (Fig 5A). Indeed, MOG induction resulted in a significant increase in the total number of T cells in the retina in comparison to naïve controls (Veh: 32 ± 8.57, MOG: 161 ± 40, n = 4/group, p<0.05; Fig 5B). Notably, CCR3 inhibition significantly reduced the number of MOG-induced T cells in the retina in comparison to untreated MOG retinas (MOG + AKST4290: 73 ± 19.21, n = 9, p<0.05; Fig 5B).

**Fig 5.**
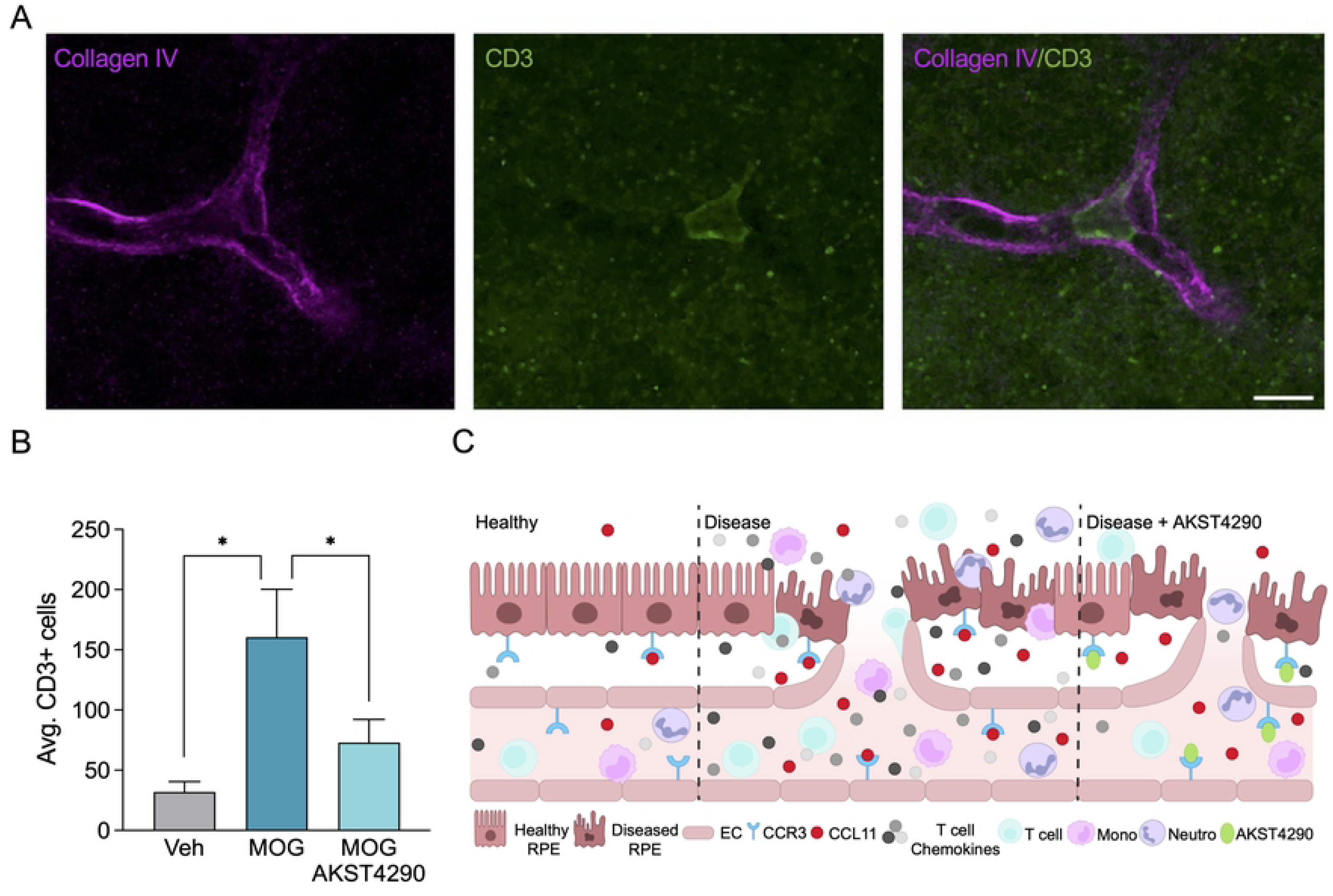
CCR3 inhibition reduces retinal T cell infiltration in a model of induced autoimmune inflammatory disease. (A) Retinal flatmount stained with collagen IV (magenta) labeling vasculature and CD3 (green) labeling T cells. (B) Quantification of CD3+ T cells in retinas treated with Veh, MOG, or MOG + AKST4290 (n=4,4,9). (C) Potential mechanism of action of CCR3 inhibition in inflammatory ocular diseases. Mean ± SEM. *p<0.05; Mann-Whitney; Scale bar: 10 mm. Veh: vehicle; MOG: myelin oligodendrocyte glycoprotein; EC: endothelial cell; Mono: monocyte; Neutro: neutrophil.

## Discussion

Circulating populations of T cells in the blood are known to be dysregulated in AMD patients [26]. In addition, studies have shown T cells are present in ocular tissues in AMD animal models and patients [26,27]. However, the precise function of lymphocytes in driving pathologies in AMD remains poorly understood [26,27]. Interestingly, peripheral T cells expressing CCR3 and the primary ligand for CCR3, CCL11 or eotaxin, are present in CNV lesions of nAMD animal models and patient tissue samples [17,18,28]. Furthermore, increased levels of CCL11 were reported in the aqueous humor of nAMD patients as well as in the ocular tissue of dry AMD patients [29–31]. Our data show that NaIO_3_ injury results in a significant increase in ocular levels of CCL11 and we hypothesized that increased levels of CCL11 in the eye could act as an attractant source for peripheral immune cells. Indeed, we discovered that CCR3 inhibition in the NaIO_3_ model significantly reduced disease-induced increase in T cell subtypes in the RPE/choroid (Figs 3 and 4). Furthermore, we validated the impact of CCR3 inhibition on the recruitment of T cells in the MOG model (Fig 5). These results are consistent with the observed effects of CCR3 inhibition by AKST4290 in reducing the number of T cells in the brain in the MOG model [23].

Infiltrating immune cells release cytokines in response to the inflammatory environment in the diseased tissue which in turn further exacerbates local inflammation thereby creating a feedback loop that perpetuates the inflammatory cycle [19,32]. Previous *in vitro* studies have shown that activated T cells induce RPE cells to secrete multiple cytokines including CCL5 and CXCL9 [33]. CCL5 and CXCL9 cytokines are implicated in multiple pathological processes including inflammation [34–38]. Importantly, their function as potent attractants for T cells, including CD4+ T cells, is well established. Thus, these chemokines can contribute to a cycle of damage propagating inflammation [32,37]. The data presented here show that NaIO_3_ injection causes an elevation of CCL5 and CXCL9 concentrations in the eye which is significantly reduced following inhibition of CCR3 with AKST4290 suggesting CCR3 inhibition plays an anti-inflammatory role (Fig 3).

Our findings indicate that there are two potential sites of AKST4290 action in namely the RPE/choroid and circulating immune cells. Due to the oral systemic dosing route of AKST4290 a limitation of this study is we cannot parse the effect of CCR3 inhibition on peripheral immune cells from the effect of CCR3 inhibition on cells in the RPE/choroid. Although beyond the scope of the current study, future studies using cell-type specific CCR3 knockout mice and site-specific delivery of CCR3 inhibition would help disambiguate the systemic versus local contribution of CCR3 to ocular inflammation.

Furthermore, this approach would also allow examination of the direct effect of CCR3 inhibition on RPE cell survival and function under inflammatory stimuli.

Interestingly, a recent study by Rege et al. reported that multiple CCR3 ligands including CCL11 increase in the mouse brain with age; and inhibition of CCR3 with orally-dosed AKST4290 was demonstrated to reduce the age-dependent increase in T cells in the brain [23,39]. They confirmed that peripheral T cells do indeed express CCR3 and went on to reveal that CCR3 expression in the brain was limited to a very small proportion of microglia. Moreover, AKST4290 was reported to have marginal brain penetrance due to polar surface area and high molecular weight. These findings led the authors to postulate a peripheral site of action for the observed effects in the brain with CCR3 inhibition which differs from our findings of multiple sites of action in ocular diseases. Together, these data further underscore the necessity to study the function of CCR3 in an organ and context dependent manner to better understand the mechanisms of action of CCR3 inhibition in different diseases.

Curiously, we also observed a profound impact of CCR3 inhibition on infiltration of other immune cell types that do not express CCR3 (Fig 3). Under activated conditions both neutrophils and monocytes can be recruited by CCL5 [40,41]. Therefore, the reduction in neutrophils and monocytes with CCR3 inhibition in NaIO_3_ damaged RPE/choroid could potentially be due to the reduction of chemokines such as CCL5. Another possibility is that the change in different immune populations reflects the crosstalk between different types of immune cells [42–44]. For example, IL-17 secreted by activated CD4+ T cells functions in a pro-inflammatory manner to recruit neutrophils [45]. Ultimately, our findings add to the growing body of evidence highlighting the complex interplay between different immune populations and chemokines in disease.

Overall, the findings in this study extend the immunomodulatory role of CCR3 in the eye. We propose a model (Fig 5C) in which disease associated inflammation supports homing of peripheral immune cells to sites of damage. The interaction of infiltrating cells and the local milieu drives pathogenesis. In this scenario, CCR3 inhibition could act as a break for persistent inflammation and therapeutic target.

## Conclusion

In conclusion, we elucidated an anti-inflammatory role for CCR3 inhibition in two models of acute ocular inflammation. Our data revealed that CCR3 inhibition with AKST4290 reduces the number of infiltrating immune cells and concomitantly reduces inflammatory chemokines in the diseased eye. Together, these findings strengthen CCR3’s position as an interesting molecular node of immune modulation for ocular inflammatory diseases such as nAMD.

## Acknowledgements

The authors thank J. Masumi and B. Higgins for technical assistance, B. Von Melchert for vivarium support, and E. Newman, S. Garg, and E. Jeffords for critical discussions.

## Supporting information

**S1 Fig. RPE degeneration in the NaIO_3_ model.**

RPE flatmount stained with phalloidin (white) and DAPI (blue) showing RPE degeneration (dotted circle) at 3 days post single IV injection of (A) vehicle or (B) 20mg/kg NaIO_3_. Right panel is 200 magnification of corresponding left panel white box.

**S2 Fig. CCR3 inhibition effects on cytokines in the eye in the NaIO_3_ model.**

n=13-30 mice/group. Mean ± SEM. *p<0.05, **p<0.01, ****p<0.0001; nested linear model followed by Tukey’s posthoc analysis. ns: not significant.

**S3 Table. CCR3 expression on RPE and endothelial cells.**

Proportion of RPE and EC expressing CCR3 (n=5). EC: endothelial cells; RPE: retinal pigment epithelial cells.

**S4 Table. QWBA detection of AKST4290 in the eye.**

Quantification of [14C]-AKST4290 levels after a single PO dose of 10mg (100 µCi/kg) in male and female whole eye and uveal tract (n=1/timepoint). QWBA: quantitative whole-body autoradiography; PO: oral gavage; LLOQ: lower limit of quantification.

**S5 Table. LC-MS/MS detection of AKST4290 in eye compartments.**

Quantification of AKST4290 levels in eye sub compartments after a single 30 mg/kg PO dose (n=1 at timepoint 0, n=4-5/all other timepoints). LC-MS/MS: liquid chromatography with tandem mass spectrometry; PO: oral gavage; RPE: retinal pigment epithelial cells.

**S6 Table. Radius ratio measurement of RPE damage.**

Quantification of radius ratio (average radial RPE damage) in RPE flatmounts of mice treated with vehicle, NaIO_3_, or NaIO_3_ + AKST4290. n=15-17 mice/group. RPE: retinal pigment epithelial cells.

**S7 Table. Inflammatory chemokine levels in the RPE.**

RPE chemokine levels in the RPE of mice treated with vehicle, NaIO_3_, or NaIO_3_ + AKST4290. n=13-30 mice/group. RPE: retinal pigment epithelial cells.

**S8 Table. Quantification of immune cell populations in mice treated with Vehicle, NaIO_3_, or NaIO_3_ + AKST4290.**

Immune cell counts in the RPE. n=12 mice/group. RPE: retinal pigment epithelial cells.

**S9 Table. Quantification of CD3+ T cells in retinas treated with Veh, MOG, or MOG + AKST4290.**

T cell quantification in the retina. n=4-9 mice/group. Veh: vehicle; MOG: myelin oligodendrocyte glycoprotein.

## References

1. Tan W, Zou J, Yoshida S, Jiang B, Zhou Y. The Role of Inflammation in Age-Related Macular Degeneration. Int J Biol Sci. 2020;16: 2989–3001. doi:10.7150/ijbs.49890

2. Kauppinen A, Paterno JJ, Blasiak J, Salminen A, Kaarniranta K. Inflammation and its role in age-related macular degeneration. Cellular and Molecular Life Sciences. 2016;73: 1765–1786. doi:10.1007/s00018-016-2147-8

3. Ambati J, Fowler BJ. Mechanisms of Age-Related Macular Degeneration. Neuron. 2012;75: 26–39. doi:10.1016/j.neuron.2012.06.018

4. Cao L, Wang H, Wang F, Xu D, Liu F, Liu C. Aβ-Induced Senescent Retinal Pigment Epithelial Cells Create a Proinflammatory Microenvironment in AMD. Investigative Opthalmology & Visual Science. 2013;54: 3738. doi:10.1167/iovs.13-11612

5. Kinnunen K, Petrovski G, Moe MC, Berta A, Kaarniranta K. Molecular mechanisms of retinal pigment epithelium damage and development of age-related macular degeneration. Acta Ophthalmol. 2012;90: 299–309. doi:10.1111/j.1755-3768.2011.02179.x

6. Anderson DH, Mullins RF, Hageman GS, Johnson L V. A role for local inflammation in the formation of drusen in the aging eye. Am J Ophthalmol. 2002;134: 411–431. doi:10.1016/S0002-9394(02)01624-0

7. Flores R, Carneiro Â, Vieira M, Tenreiro S, Seabra MC. Age-Related Macular Degeneration: Pathophysiology, Management, and Future Perspectives. Ophthalmologica. 2021;244: 495–511. doi:10.1159/000517520

8. Hadziahmetovic M, Malek G. Age-Related Macular Degeneration Revisited: From Pathology and Cellular Stress to Potential Therapies. Front Cell Dev Biol. 2021;8. doi:10.3389/fcell.2020.612812

9. Liao DS, Grossi F V., El Mehdi D, Gerber MR, Brown DM, Heier JS, et al. Complement C3 Inhibitor Pegcetacoplan for Geographic Atrophy Secondary to Age-Related Macular Degeneration. Ophthalmology. 2020;127: 186–195. doi:10.1016/j.ophtha.2019.07.011

10. Ghasemi Falavarjani K, Nguyen QD. Adverse events and complications associated with intravitreal injection of anti-VEGF agents: a review of literature. Eye. 2013;27: 787–794. doi:10.1038/eye.2013.107

11. Yerramothu P. New Therapies of Neovascular AMD—Beyond Anti-VEGFs. Vision. 2018;2: 31. doi:10.3390/vision2030031

12. Patel P, Sheth V. New and Innovative Treatments for Neovascular Age-Related Macular Degeneration (nAMD). J Clin Med. 2021;10: 2436. doi:10.3390/jcm10112436

13. Stewart MW, Garg S, Newman EM, Jeffords E, Konopińska J, Jackson S, et al. Safety and Therapeutic Effects of Orally Administered AKST4290 in Newly Diagnosed Neovascular Age-Related Macular Degeneration. Retina. 2022;42: 1038–1046. doi:10.1097/IAE.0000000000003446

14. Salcedo R, Young HA, Ponce ML, Ward JM, Kleinman HK, Murphy WJ, et al. Eotaxin (CCL11) Induces In Vivo Angiogenic Responses by Human CCR3+ Endothelial Cells. The Journal of Immunology. 2001;166: 7571–7578. doi:10.4049/jimmunol.166.12.7571

15. Sokol CL, Luster AD. The Chemokine System in Innate Immunity. Cold Spring Harb Perspect Biol. 2015;7: a016303. doi:10.1101/cshperspect.a016303

16. Francis JN, Lloyd CM, Sabroe I, Durham SR, Till SJ. T lymphocytes expressing CCR3 are increased in allergic rhinitis compared with non-allergic controls and following allergen immunotherapy. Allergy. 2007;62: 59–65. doi:10.1111/j.1398-9995.2006.01253.x

17. Nagai N, Ju M, Izumi-Nagai K, Robbie SJ, Bainbridge JW, Gale DC, et al. Novel CCR3 Antagonists Are Effective Mono-and Combination Inhibitors of Choroidal Neovascular Growth and Vascular Permeability. Am J Pathol. 2015;185: 2534– 2549. doi:10.1016/j.ajpath.2015.04.029

18. Takeda A, Baffi JZ, Kleinman ME, Cho WG, Nozaki M, Yamada K, et al. CCR3 is a target for age-related macular degeneration diagnosis and therapy. Nature. 2009;460: 225–230. doi:10.1038/nature08151

19. Sallusto F, Mackay CR, Lanzavecchia A. The Role of Chemokine Receptors in Primary, Effector, and Memory Immune Responses. Annu Rev Immunol. 2000;18: 593–620. doi:10.1146/annurev.immunol.18.1.593

20. Targowski T, Jahnz-Rózyk K, Plusa T, Glodzinska-Wyszogrodzka E. Influence of age and gender on serum eotaxin concentration in healthy and allergic people. J Investig Allergol Clin Immunol. 2005;15: 277–82.

21. Bettcher BM, Fitch R, Wynn MJ, Lalli MA, Elofson J, Jastrzab L, et al. MCP-1 and eotaxin-1 selectively and negatively associate with memory in MCI and Alzheimer’s disease dementia phenotypes. Alzheimer’s & Dementia: Diagnosis, Assessment & Disease Monitoring. 2016;3: 91–97. doi:10.1016/j.dadm.2016.05.004

22. Moriguchi M, Nakamura S, Inoue Y, Nishinaka A, Nakamura M, Shimazawa M, et al. Irreversible Photoreceptors and RPE Cells Damage by Intravenous Sodium Iodate in Mice Is Related to Macrophage Accumulation. Invest Ophthalmol Vis Sci. 2018;59: 3476–3487. doi:10.1167/iovs.17-23532

23. Rege S V., Teichert A, Masumi J, Dhande OS, Harish R, Higgins BW, et al. CCR3 plays a role in murine age-related cognitive changes and T-cell infiltration into the brain. Commun Biol. 2023;6: 292. doi:10.1038/s42003-023-04665-w

24. Plaza Reyes A, Petrus-Reurer S, Padrell Sánchez S, Kumar P, Douagi I, Bartuma H, et al. Identification of cell surface markers and establishment of monolayer differentiation to retinal pigment epithelial cells. Nat Commun. 2020;11: 1609. doi:10.1038/s41467-020-15326-5

25. Constantinescu CS, Farooqi N, O’Brien K, Gran B. Experimental autoimmune encephalomyelitis (EAE) as a model for multiple sclerosis (MS). Br J Pharmacol. 2011;164: 1079–1106. doi:10.1111/j.1476-5381.2011.01302.x

26. Behnke V, Wolf A, Langmann T. The role of lymphocytes and phagocytes in age-related macular degeneration (AMD). Cellular and Molecular Life Sciences. 2020;77: 781–788. doi:10.1007/s00018-019-03419-4

27. Camelo S. Potential Sources and Roles of Adaptive Immunity in Age-Related Macular Degeneration: Shall We Rename AMD into Autoimmune Macular Disease? Autoimmune Dis. 2014;2014: 1–11. doi:10.1155/2014/532487

28. Mizutani T, Ashikari M, Tokoro M, Nozaki M, Ogura Y. Suppression of Laser-Induced Choroidal Neovascularization by a CCR3 Antagonist. Invest Ophthalmol Vis Sci. 2013;54: 1564–1572. doi:10.1167/iovs.11-9095

29. Sato T, Takeuchi M, Karasawa Y, Enoki T, Ito M. Intraocular inflammatory cytokines in patients with neovascular age-related macular degeneration before and after initiation of intravitreal injection of anti-VEGF inhibitor. Sci Rep. 2018;8: 1098. doi:10.1038/s41598-018-19594-6

30. Sun T, Wei Q, Gao P, Zhang Y, Peng Q. Cytokine and Chemokine Profile Changes in Patients with Neovascular Age-Related Macular Degeneration After Intravitreal Ranibizumab Injection for Choroidal Neovascularization. Drug Des Devel Ther. 2021;Volume 15: 2457–2467. doi:10.2147/DDDT.S307657

31. Mo FM, Proia AD, Johnson WH, Cyr D, Lashkari K. Interferon γ–Inducible Protein-10 (IP-10) and Eotaxin as Biomarkers in Age-Related Macular Degeneration. Investigative Opthalmology & Visual Science. 2010;51: 4226. doi:10.1167/iovs.09-3910

32. Rhee MK, Mah FS. Inflammation in Dry Eye Disease. Ophthalmology. 2017;124: S14–S19. doi:10.1016/j.ophtha.2017.08.029

33. Juel HB, Faber C, Udsen MS, Folkersen L, Nissen MH. Chemokine Expression in Retinal Pigment Epithelial ARPE-19 Cells in Response to Coculture with Activated T Cells. Invest Ophthalmol Vis Sci. 2012;53: 8472–8480. doi:10.1167/iovs.12-9963

34. Lacotte S, Brun S, Muller S, Dumortier H. CXCR3, Inflammation, and Autoimmune Diseases. Ann N Y Acad Sci. 2009;1173: 310–317. doi:10.1111/j.1749-6632.2009.04813.x

35. Detrick B, Hooks JJ. Immune regulation in the retina. Immunol Res. 2010;47: 153–161. doi:10.1007/s12026-009-8146-1

36. Müller M, Carter S, Hofer MJ, Campbell IL. Review: The chemokine receptor CXCR3 and its ligands CXCL9, CXCL10 and CXCL11 in neuroimmunity - a tale of conflict and conundrum. Neuropathol Appl Neurobiol.2010;36: 368–387. doi:10.1111/j.1365-2990.2010.01089.x

37. Marques RE, Guabiraba R, Russo RC, Teixeira MM. Targeting CCL5 in inflammation. Expert Opin Ther Targets. 2013;17: 1439–1460. doi:10.1517/14728222.2013.837886

38. Zeng Z, Lan T, Wei Y, Wei X. CCL5/CCR5 axis in human diseases and related treatments. Genes Dis. 2022;9: 12–27. doi:10.1016/j.gendis.2021.08.004

39. Baruch K, Ron-Harel N, Gal H, Deczkowska A, Shifrut E, Ndifon W, et al. CNS-specific immunity at the choroid plexus shifts toward destructive Th2 inflammation in brain aging. Proceedings of the National Academy of Sciences. 2013;110: 2264–2269. doi:10.1073/pnas.1211270110

40. Hwaiz R, Rahman M, Syk I, Zhang E, Thorlacius H. Rac1-dependent secretion of platelet-derived CCL5 regulates neutrophil recruitment via activation of alveolar macrophages in septic lung injury. J Leukoc Biol. 2015;97: 975–984. doi:10.1189/jlb.4A1214-603R

41. Schall TJ, Bacon K, Toy KJ, Goeddel D V. Selective attraction of monocytes and T lymphocytes of the memory phenotype by cytokine RANTES. Nature. 1990;347: 669–671. doi:10.1038/347669a0

42. Stepp MA, Menko AS. Immune responses to injury and their links to eye disease. Translational Research. 2021;236: 52–71. doi:10.1016/j.trsl.2021.05.005

43. Goverman J. Autoimmune T cell responses in the central nervous system. Nat Rev Immunol. 2009;9: 393–407. doi:10.1038/nri2550

44. Ransohoff RM, Brown MA. Innate immunity in the central nervous system. Journal of Clinical Investigation. 2012;122: 1164–1171. doi:10.1172/JCI58644

45. Laan M, Cui Z-H, Hoshino H, Lötvall J, Sjöstrand M, Gruenert DC, et al. Neutrophil Recruitment by Human IL-17 Via C-X-C Chemokine Release in the Airways. The Journal of Immunology. 1999;162: 2347. Available: http://www.jimmunol.org/content/162/4/2347.abstract

